# Micropillar enhanced FRET-CRISPR biosensor for nucleic acid detection

**DOI:** 10.1101/2023.08.23.554533

**Authors:** Mengdi Bao, Stephen J. Dollery, FNU Yuqing, Gregory J. Tobin, Ke Du

## Abstract

CRISPR technology has gained widespread adoption for pathogen detection due to its exceptional sensitivity and specificity. Although recent studies have investigated the potential of high-aspect-ratio microstructures in enhancing biochemical applications, their application in CRISPR-based detection has been relatively rare. In this study, we developed a FRET-based biosensor in combination with high-aspect-ratio microstructures and Cas12a-mediated trans-cleavage for detecting HPV 16 DNA fragments. Remarkably, our results show that micropillars with higher density exhibit superior molecular binding capabilities, leading to a tenfold increase in detection sensitivity. Furthermore, we investigated the effectiveness of two surface chemical treatment methods for enhancing the developed FRET assay. A simple and effective approach was also developed to mitigate bubble generation in microfluidic devices, a crucial issue in biochemical reactions within such devices. Overall, this work introduces a novel approach using micropillars for CRISPR-based viral detection and provides valuable insights into optimizing biochemical reactions within microfluidic devices.

## Introduction

Clustered Regularly Interspaced Short Palindromic Repeats system (CRISPR) has emerged as a groundbreaking technology with vast potential in numerous biomedical applications^1, 2^, particularly in the detection of nucleic acid-based infectious diseases.^3, 4^ The fundamental principle of CRISPR-based detection involves the design of CRISPR-RNAs (crRNAs) that possess sequences complementary to specific viral targets.^5^ This design allows the associated Cas proteins to accurately locate and cleave these target sequences with excellent specificity.^6^ Recent discoveries have shed light on the trans-cleavage activity exhibited by Cas12a and Cas13 proteins, making them widely adopted in viral detection applications.^7, 8^ These proteins demonstrate the remarkable ability to cleave non-targeted single-stranded nucleic acids once they have identified the targets. By incorporating fluorophore-quencher probes, activated Cas proteins non-specifically cleave these probes, thereby amplifying the detection signals and showcasing a high level of sensitivity.^9, 10^ Additionally, the CRISPR-Cas detection offers the advantage of being performed isothermally, providing an excellent alternative to reverse transcription-polymerase chain reaction (RT-PCR) detection.

In recent years, the use of high-aspect-ratio micro/nanostructures in biochemical applications has sparked a growing interest.^11, 12^ These structures provide more binding sites for molecules and offer morphological features that promote effective interactions with surface probes during biochemical reactions.^13, 14^ As an example, zinc oxide (ZnO) nanorods were integrated as the working electrode in an electrochemical biosensor.^15^ These nanostructures offer an extended surface area for immobilizing antibodies, resulting in significantly lower limits of detection compared to traditional assays. In a separate work, ZnO served as a scaffold for capturing avian influenza virus.^16^ When incorporated with the gold-nanoparticle colorimetric reaction, the ZnO-based detection platform demonstrated one order of magnitude improvement sensitivity over conventional fluorescence-based ELISA. Similarity, in an example of microstructures, Movilli et al. introduced densely structured micropillars for electrochemical DNA detection.^17^ The high-aspect-ratio micropillar array displayed up to one order of magnitude higher sensitivity compared to conventional flat substrates. Likewise, Bandaru et al. developed a high-throughput micropillar and microwell array, leading to a remarkable improvement in the capture efficiency of biomarkers secreted by cancer cells.^18^

In our previous research, we successfully demonstrated the enhanced molecular binding achieved using micropillars fabricated through photolithography.^19^ Additionally, we highlighted the versatility of the micropillar arrays as effective platforms for CRISPR-based nucleic acid detection, showing that a 40% higher probe binding load is achieved with the extend surface.^19^ However, the aspect ratio of the micropillars was limited at ∼1.2:1 due to fabrication constrains. In addition, the quantitative correlation between micropillar density and binding capacity remained unexplored. Therefore, in this study, we aimed to address two crucial knowledge gaps: (1) investigating the impact of density variations on molecular binding capacities, and (2) establishing a direct correlation between micropillar density and detection sensitivity, thereby evaluating their potential for enhancing CRISPR-based nucleic acid detection. To achieve these objectives, we adopted a laser micromachining approach to fabricate higher-aspect-ratio (height: 300 µm, top diameter: 25 µm) polydimethylsiloxane (PDMS) micropillars, rather than using the conventional photolithography method. We introduced a Förster resonance energy transfer (FRET)-based biosensor, combined with Cas12a-mediated trans-cleavage for the detection of Human Papillomavirus (HPV) targets. Quantum dots (QDs) were chosen as the FRET donor, while single-stranded DNA (ssDNA) with an Iowa RQ linker (quencher probes) acted as the acceptor, quenching the fluorescence intensity of QDs. Upon the introduction of the target, activated Cas12a degraded the ssDNA quencher probes, resulting in QDs emitting high fluorescence signals in the channel.

Our results demonstrated that micropillars with higher density exhibited enhanced molecular binding and improved detection sensitivity, establishing a clear connection between micropillar density and their performance in viral detection. Additionally, we explored two surface chemical treatment methods to determine their effectiveness for the developed FRET assay. Furthermore, we presented a simple and effective approach to reduce bubble generation in microfluidic devices, addressing a crucial issue in biochemical reactions within such devices. This study not only presents a novel research approach utilizing micropillars for CRISPR-based viral detection, but also offers valuable insights into improving biochemical reactions within microfluidic devices.

## Results

In this study, we developed a FRET-based biosensor combined with high-aspect-ratio microstructures and Cas12a-mediated trans-cleavage for the detection of HPV 16 DNA fragment. The process is depicted in **Figure 1a**. In our designed FRET assay, QDs are selected as the FRET donor, while single-stranded DNA (ssDNA) with an Iowa RQ linker serves as the acceptor to quench the fluorescence intensity of QDs. Upon binding to the target nucleic acids, Cas12a undergoes a conformational change, transforming into nonspecific single-stranded DNA nucleases. This enzymatic activity results in the cleavage of the single-stranded DNA and the subsequent release of the Iowa RQ linker into the solution. As a result, the Iowa RQ linker is unable to approach the QDs closely, allowing the QDs to emit strong fluorescence signals. In the absence of the target input, biotinylated Iowa RQ-modified ssDNA molecules are covalently linked to streptavidin-modified QDs, resulting in the quenching of the QD’s fluorescence intensity. This FRET assay allows for the translation of CRISPR-target identification and Cas12a-mediated trans-cleavage into a fluorescence signal difference between the channel with and without target input. **Figure 1b** illustrates the measurement process, in which fluorescence microscope is employed to capture high-resolution images of the 5 cm x 5 cm micropillar area within the channels. The acquired images were subsequently processed and analyzed using the ImageJ software, enabling the calculation of fluorescence intensity. This quantified fluorescence intensity was then utilized to determine the concentration of the target input, also providing information for the evaluation of the micropillar detection’s performance. An example illustrating the fluorescence difference between a channel with target input and a channel without target input is presented in **Figure 1c**. In the fluorescence image of the channel with target input, a distinct red color is observed on the micropillars, indicating a strong fluorescence signal. Conversely, the fluorescence image of the channel without target input exhibits a diminished red color, suggesting a weaker fluorescence signal.

**Figure 1.**
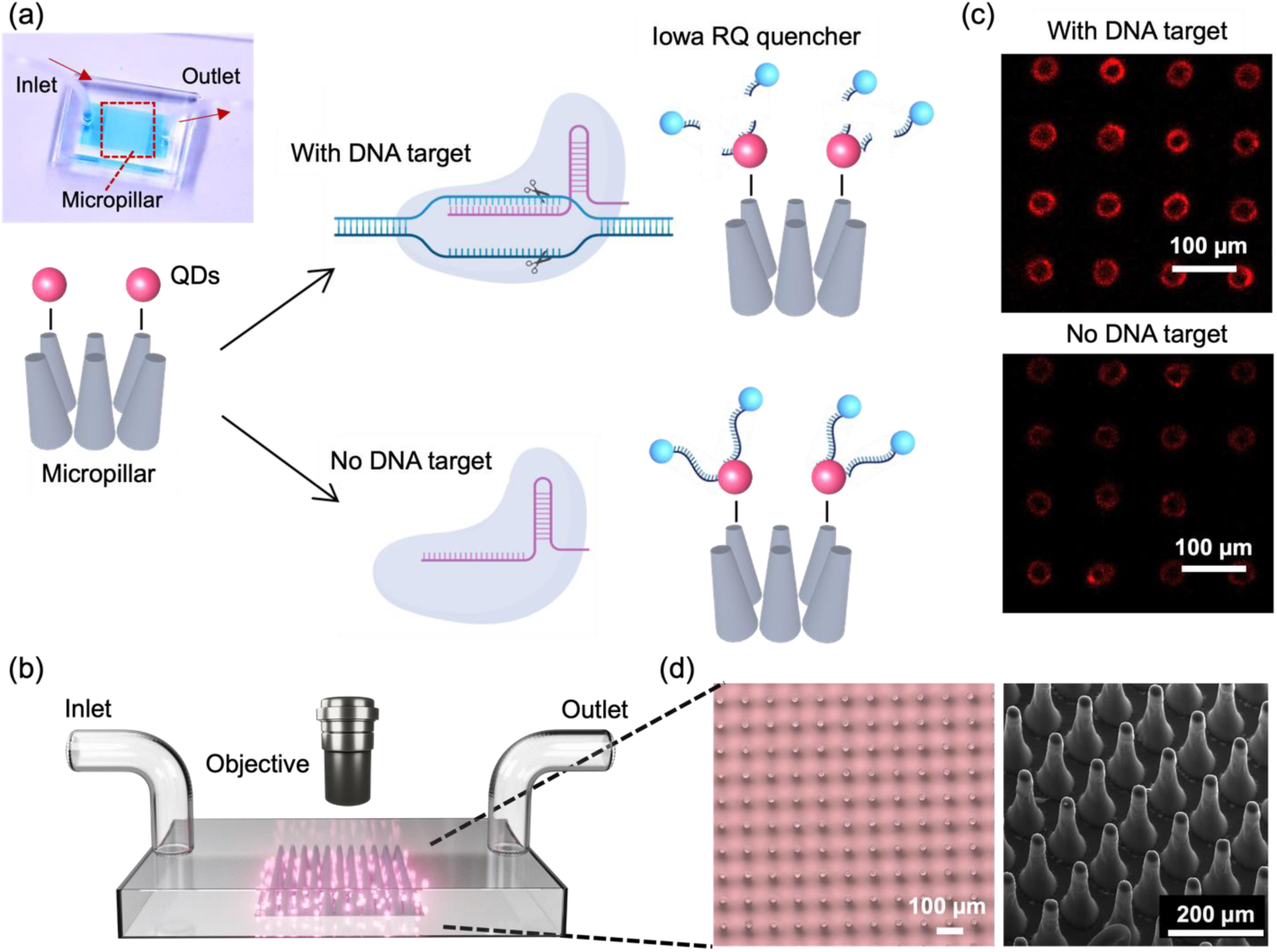
(a) Working principle of the micropillar-enhanced FRET sensor for the detection of HPV 16 fragment. Inset: photograph of the sensor filled with food dye. (b) Illustration outlining the measurement process of the sensor. (c) Fluorescent image of micropillars after reaction, in the presence (top) or absence (bottom) of the HPV 16 fragment. Scale bar: 100 µm. (d) Left: Optical micrographs of micropillars. Scale bar: 100 µm. Right: SEM image of micropillars. Scale bar: 200 µm.

To fabricate the micropillars, a laser micromachining approach was employed to create through-microholes with a diameter of 25 µm on a tungsten substrate (thickness: 300 µm).^20^ Subsequently, a mixture of PDMS and its curing agent was prepared and poured onto the tungsten substrate, forming PDMS micropillars through the process of soft lithography.^21^ The utilization of laser micromachining was preferred over traditional photolithography, as it overcomes the limitations of low-aspect-ratio and enables the production of micropillars with a taper angle. This taper angle (∼3°) proved advantageous for facilitating the observation of conjugated QDs on the side walls of the micropillars through a top-down view in microscope. **Figure 1d** presents the microscope and SEM images of the micropillar array, providing visual evidence of the uniform shape, consistent spacing, and presence of a taper angle in each micropillar.

Having successfully fabricated well-defined micropillars, we proceeded to investigate the relationship between micropillar density and its molecular binding capacity. To examine this, we prepared micropillar arrays with three different pitch distances (109 µm, 147 µm, and 197 µm) and conjugated them with QDs. We evaluated the binding capacity by comparing the fluorescence intensity of the conjugated QDs, and the result is presented in **Figure 2a**. Remarkably, the micropillars with a pitch distance of 109 µm exhibits the highest fluorescence signals, approximately five times greater than those observed for the 197 µm micropillars. The micropillars with a center-to-center distance of 147 µm shows the second-highest fluorescence signals, approximately three times greater than those of the 197 µm micropillars. The finding demonstrates a positive correlation between micropillar density and molecular binding capacity, indicating that higher density micropillars have a greater capacity to bind molecules. To visually validate these results, **Figure 2b** presents a fluorescence image of the micropillar array with different densities, further emphasizing that micropillars with higher density possess enhanced binding capability, allowing for increased QD conjugation.

**Figure 2.**
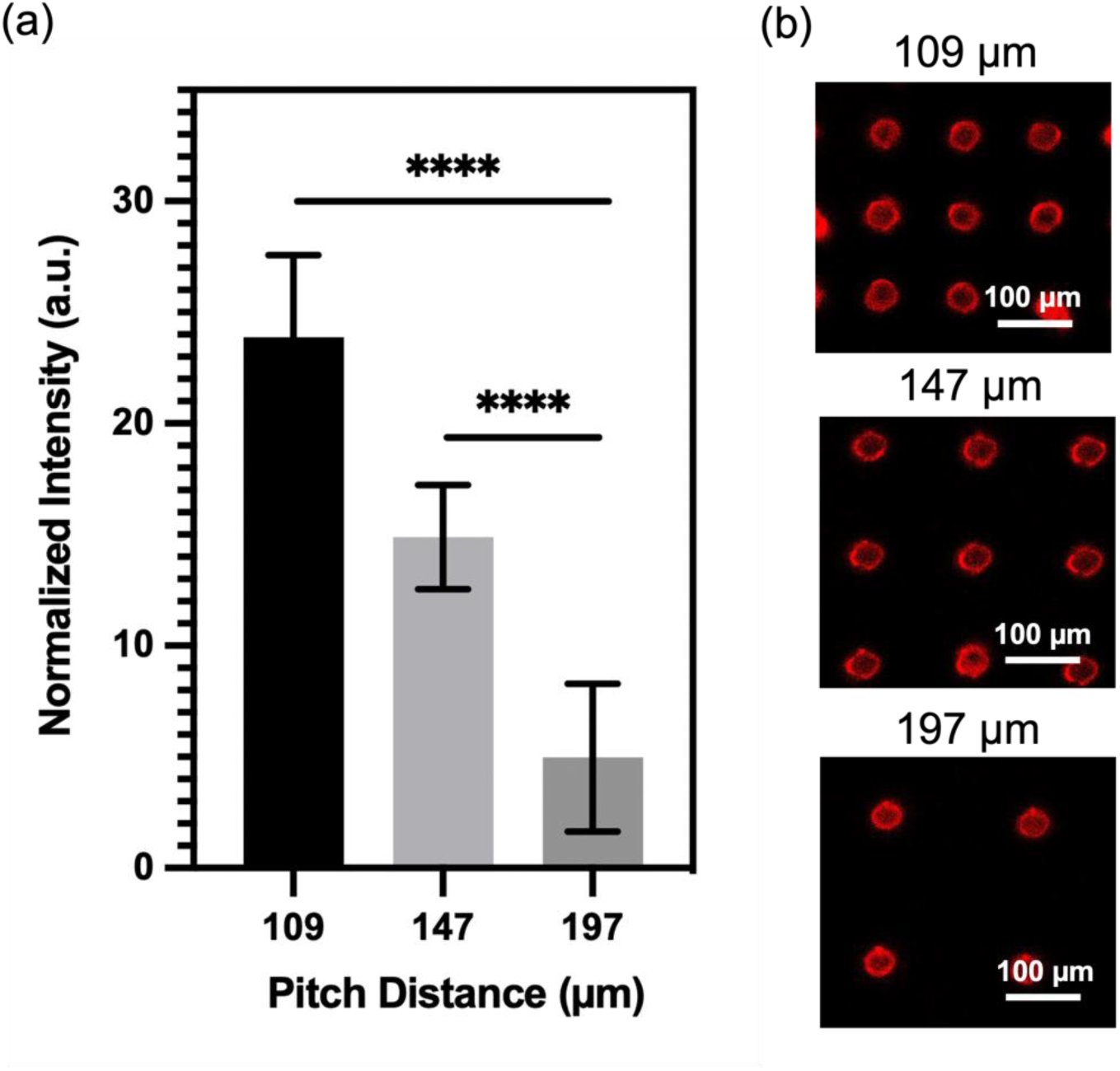
Micropillar density/molecular binding capacity relationship. (a) Normalized fluorescence signals of micropillars conjugated with QDs at different pitch distances (109 µm, 147 µm, 197 µm). Statistical analysis was conducted using unpaired t-test analysis (ns = p > 0.05; * = 0.01 < P⩽0.05; ** = 0.05 < P⩽0.05; *** = 0.01 < P⩽0.001; **** = P ⩽ 0.0001). (b) Fluorescent image of micropillars after conjugation with QDs. Micropillars tested at different pitch distances (109 µm, 147 µm, 197 µm).

As previously mentioned, the FRET assay developed in this study employed QDs as the donor and ssDNA with Iowa RQ linker as the acceptor. A crucial requirement for a successful FRET reaction is a spectral overlap between the donor’s emission and the acceptor’s absorption.^22, 23^ **Figure 3a** illustrates an overlap between the absorbance spectra of RQ and the emission spectra of QDs at 605 nm, confirming their suitability as an optimal pair for the FRET assay. Typically, in previous studies, PDMS surfaces were treated with APTMS and glutaraldehyde (GA) sequentially to immobilize streptavidin.^24, 25^ However, we observed that the reaction products of APTMS and GA could generate a Cy3 analogue with strong auto-fluorescence signals^26^, which overlaps with the absorbance spectra of the quencher probes (**Figure 3b**) and could potentially interfere with the quenching effect. To address the issue of autofluorescence, an alternative method utilizing only APTMS for QD conjugation onto PDMS surface was proposed. Two identical channels were subjected to different treatments: one with APTMS alone, and the other with a combination of APTMS and GA. Following the treatments, QDs and quencher probes were immobilized onto the channels, and the fluorescence intensity was measured before and after the probe conjugation step. The results of the experiment, as shown in **Figure 3c**, reveals significant differences in the fluorescence intensity between the two treatment approaches. The channel treated with APTMS alone exhibits a substantial reduction in QD fluorescence signal after the immobilization of the quencher probes. In contrast, the channel treated with APTMS and GA shows only a slight decrease in fluorescence intensity following the probe conjugation. The findings suggest that using APTMS alone for QD conjugation is more effective in minimizing autofluorescence and facilitating an effective FRET reaction, as compared to the APTMS and GA combination.

**Figure 3.**
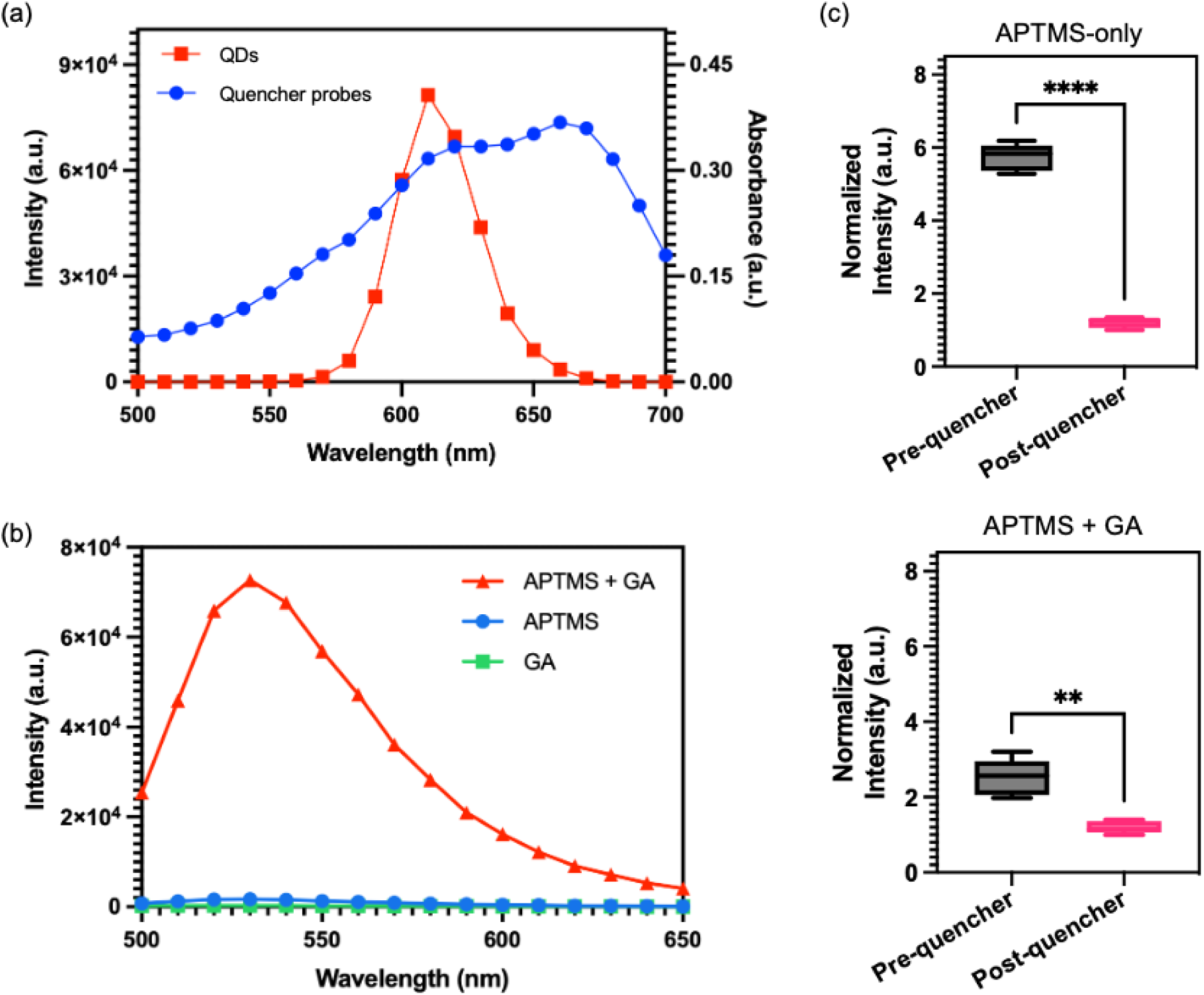
Validation of PDMS surface treatment process. (a) Emission spectra of QDs excited at 450 nm (red) and absorption spectra of quencher probes (blue). (b) Emission spectra of the mixture (red) of aminopropyltrimethoxysilane (APTMS) and glutaraldehyde (GA), APTMS (blue), and glutaraldehyde (green). (c) Normalized fluorescence signals of QDs-coated micropillars pre- and post-conjugation with quencher probes. (Top) Micropillars treated with APTMS, QDs, and quencher probes. (Bottom) Micropillars treated with APTMS, GA, QDs, and quencher probes. Statistical analysis was performed using unpaired t-test analysis (ns = p > 0.05; * = 0.01 < P⩽0.05;** = 0.05 < P⩽0.05; *** = 0.01 < P⩽0.001; **** = P ⩽ 0.0001).

To evaluate the reliability of the proposed surface modification approach, we conducted an experiment involving different concentrations of QDs conjugated onto PDMS surface. The fluorescence intensities of the PDMS were measured both before and after treatment with quencher probes to assess the effectiveness of the quenching process. **Figure 4** illustrates the results of this evaluation. Identical micropillar channels (pitch distance: 109 µm) were treated with QD concentrations ranging from 1 to 125 nM, followed by treated with 43 µM of quencher probes. Remarkably, all the QD concentrations, including 1 nM, 5 nM, 25 nM, and 125 nM, are effectively quenched by the quencher probes. The PDMS treated with 1 nM of QD exhibits the lowest difference in fluorescence intensity before and after quencher probes conjugation. This observation can be attributed to the relatively low concentration of QDs, which may result in lower fluorescence signals. Nonetheless, even at this low concentration, quencher probes are still able to effectively quench the fluorescence of QDs. This highlights the versatility of our surface modification approach in facilitating efficient quenching across a wide range of QD concentrations.

**Figure 4.**
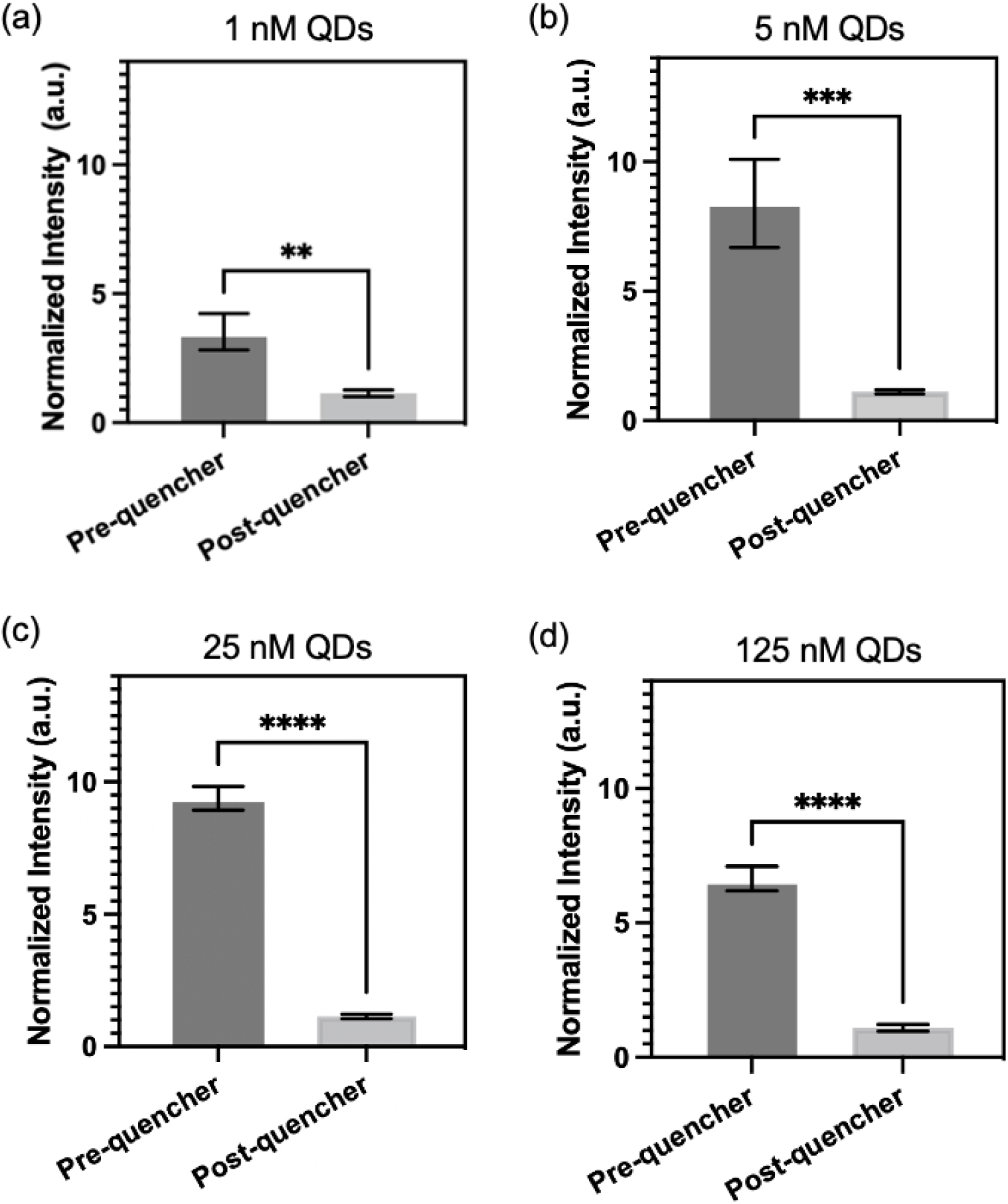
Micropillars treated with different concentrations of QDs: (a) 1 nM, (b) 5 nM, (c) 25 nM, and (d) 125 nM. Normalized fluorescence signals of micropillars measured before and after conjugation with quencher probes. Statistical analysis was performed using unpaired t-test analysis (ns = p > 0.05; * = 0.01 < P⩽0.05; ** = 0.05 < P⩽0.05; *** = 0.01 < P⩽0.001; **** = P ⩽ 0.0001).

To assess the ability of CRISPR to identify HPV 16 fragment, CRISPR-Cas12a reactions were performed in microtubes with fluorophore-quencher probes as substrates. In the presence of the target, Cas12a would undergo a transformation into a nuclease and cleave these substrates, resulting in the release of the fluorophore and generating high fluorescence signals in the solution. For systematic evaluation, six reactions labeled as #A to #F were conducted, each containing different combinations of components (**Table inside Figure 5**). The presence or absence of a specific component in each reaction was indicated by the symbols "+" and "-", respectively. The reaction was incubated at 37°C for 1 h. The results presented in **Figure 5** demonstrate that the normalized intensity of reaction #A, which contains all the necessary components, is significantly higher compared to the other reactions where certain components were lacking. This finding strongly suggests that the CRISPR-Cas system can effectively identify the HPV 16 fragment.

**Figure 5.**
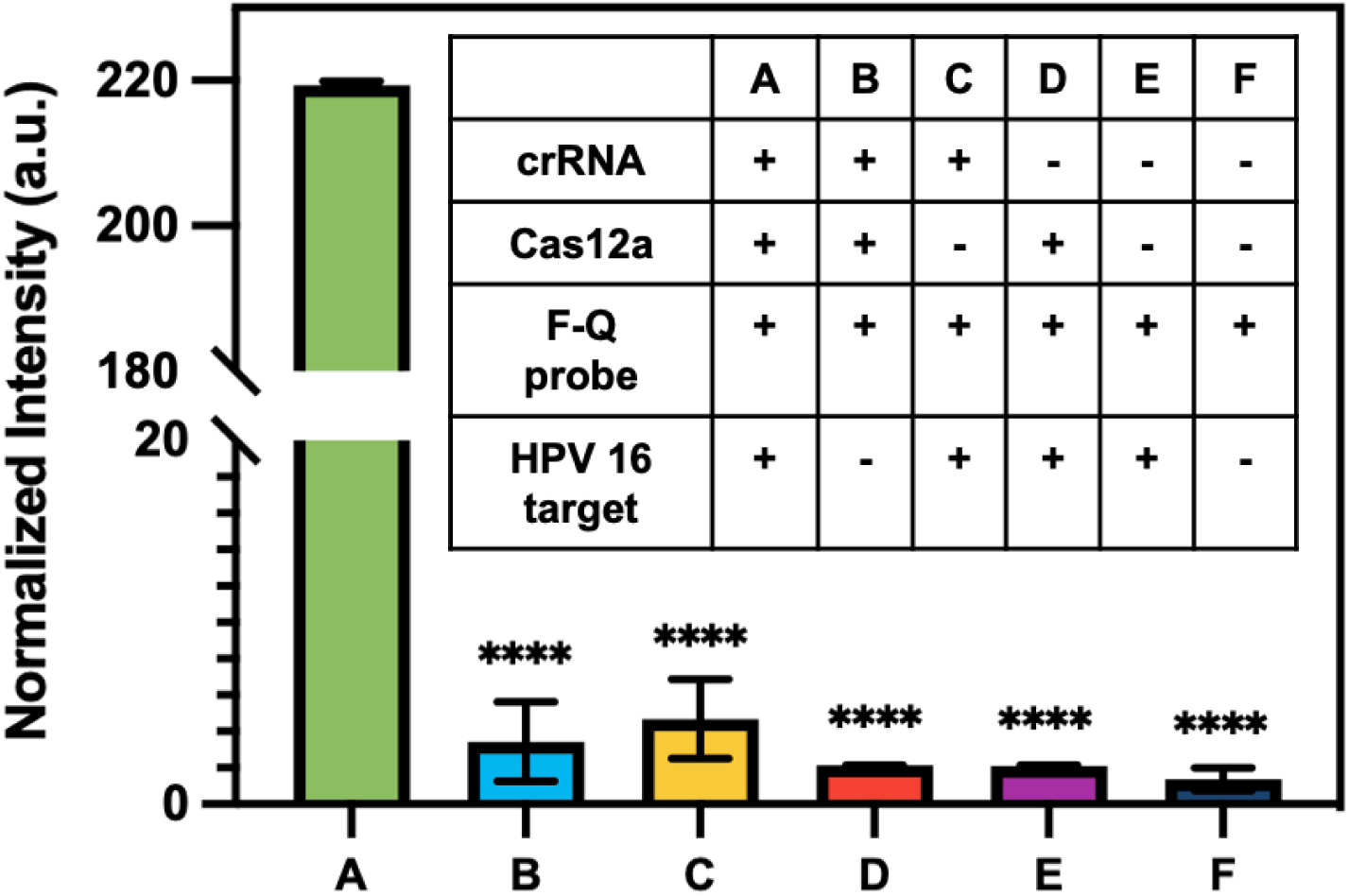
Evaluation of crRNA designed for the detection of HPV 16 fragment using fluorophore-quencher (F-Q) probes as substrates. Normalized fluorescence signals of the endpoint reactions. Statistical analysis was conducted using unpaired t-test analysis (ns = p > 0.05; * = 0.01 < P⩽0.05; ** = 0.05 < P⩽0.05; *** = 0.01 < P⩽0.001; **** = P ⩽ 0.0001). Inset: six evaluation reactions demonstrating different combinations of components. Reaction #A includes all components required for the CRISPR assay, while other reactions lack specific components.

Next, an experiment was carried out to determine the optimal concentration of quencher probes in the FRET-CRISPR detection system. The objective was to identify the concentration that would yield the most significant fluorescence difference between positive sample (with HPV 16 input) and negative control (no input). The results, depicted in **Figure 6**, reveals that the use of 30 µM quencher probes generates the highest fluorescence difference. At a concentration of 16.7 µM, the low concentration of quencher probes proves insufficient to effectively quench the fluorescence intensity of the QDs. As a result, the fluorescence difference between the positive and negative samples is not significant. When using 43 µM quencher probes, an excessive amount is present, which is not fully degraded by Cas in the positive sample. Consequently, no significant fluorescence readouts are observed in the channel. Based on this finding, a concentration of 30 µM was deemed optimal for the quencher probes in the subsequent experiments.

**Figure 6.**
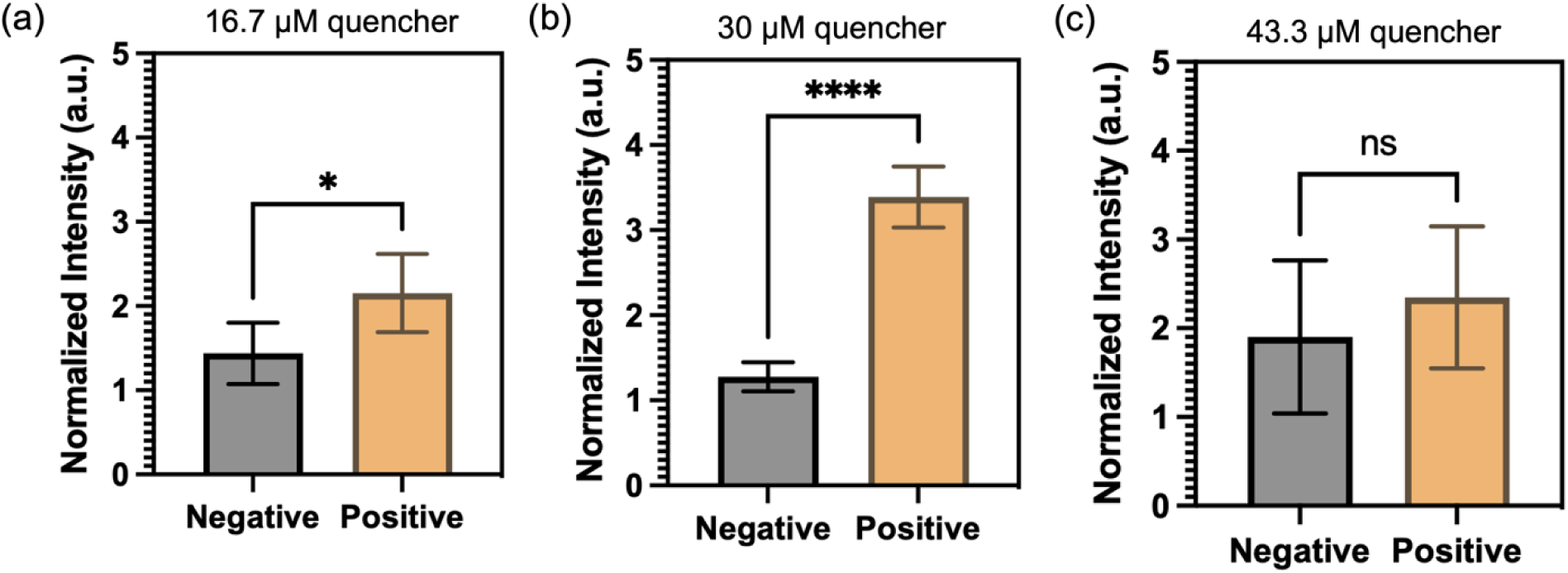
Optimization of CRISPR-Cas12a detection with varying concentrations of quencher probe: (a) 16.7 µM, (b) 30 µM, and (c) 43 µM. Normalized intensities of micropillar with the input of 10 ng/µL of HPV 16 fragment (positive) and blank (negative).

To evaluate the potential enhancement of detection sensitivity using micropillars with higher density, two types of micropillars were employed: one with a pitch distance of 197 µm and the other with a pitch distance of 109 µm. The FRET-CRISPR reaction was conducted using serially diluted HPV 16 fragments, and the assay’s sensitivity was assessed based on the fluorescence intensity of the micropillar arrays. The results, illustrated in **Figure 7a**, indicate that when using the micropillar with a pitch distance of 197 µm, the surface intensities of the target input at 10 and 1 ng/µL are higher compared to those at 0.001, 0.01, and 0.1 ng/µL. This suggests that the detection limit of the micropillars with a pitch distance of 197 µm is 1 ng/µL. **Figure 7b** demonstrates a linear decrease in intensities of 109 µm with concentrations of 0.1, 1, and 10 ng/µL, all of which are higher than those of 0.01 and 0.001 ng/µL HPV 16 target. This indicates that the micropillar with a pitch distance of 109 µm has a lower detection limit of 0.1 ng/µL. The sensitivity of the micropillars with higher density (pitch distance: 109 µm) is higher than that of the micropillars with lower density (pitch distance: 197 µm). This highlights the enhanced effectiveness of utilizing high-aspect-ratio micropillars in achieving improved detection sensitivity.

**Figure 7.**
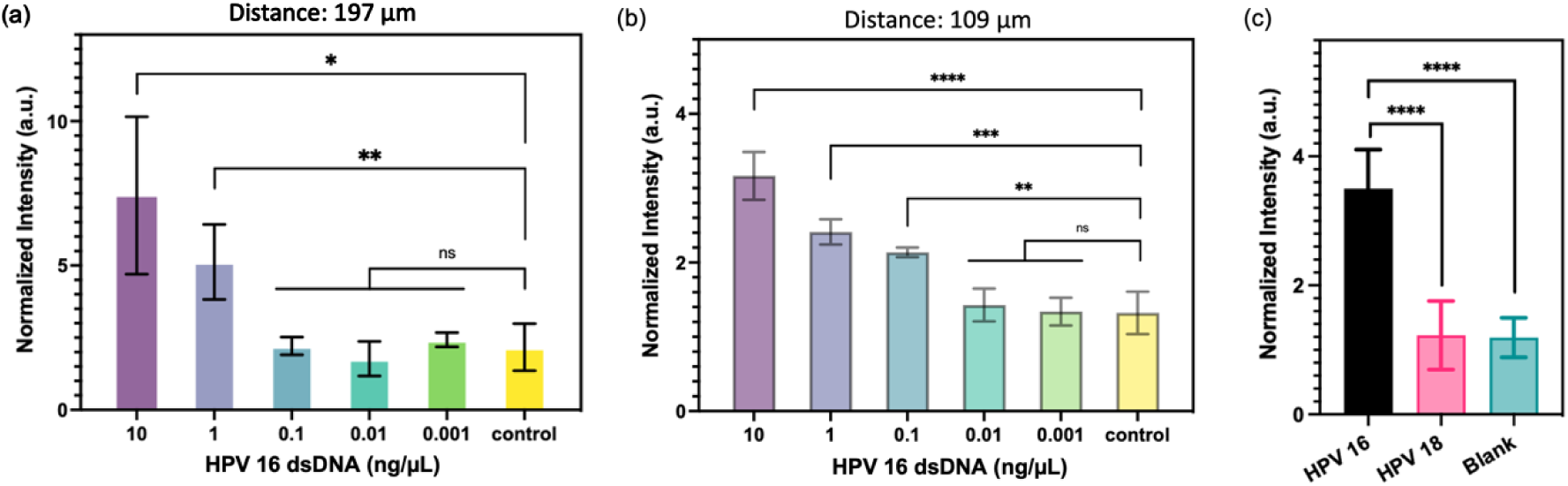
Analytical assessment of micropillars with (a) 197 µm and (b) 109 µm pitch distance for HPV 16 target detection. Normalized intensities of micropillars after CRISPR reactions are presented. (c) Normalized fluorescence signals of micropillars with the input of HPV 16 (black), HPV 18 (pink), and blank (green). Statistical analyses were conducted using unpaired t-test analysis (ns = p > 0.05; * = 0.01 < P ⩽ 0.05; ** = 0.05 < P⩽0.05; *** = 0.01 < P⩽0.001; **** = P ⩽0.0001).

As the micropillars with a pitch distance of 109 µm demonstrated better sensitivity, we proceeded to evaluate the detection specificity using these higher density micropillars. To assess specificity, we selected the HPV 18 fragment for testing, as HPV 16 and HPV 18 share significant sequence similarities. As depicted in **Figure 7c**, the channel’s intensity with HPV 16 fragment input is higher than that of HPV 18 and the blank sample. This observation indicates that the FRET-CRISPR system exhibits high detection specificity, as it can effectively distinguish between fragments with sequence similarities, such as HPV 16 and HPV 18.

## Discussion

This study investigates the potential of periodic micropillar arrays, fabricated using laser micromachining and soft lithography techniques, as effective platforms for detecting viral nucleic acids. The arrays, with their uniform micropattern, offer advantages such as consistent molecular binding and robustness during biochemical experiments, distinguishing them from random micro-and nanostructures.^27, 28^ Previous research has already demonstrated the superior molecular binding capacity of micropillars compared to planar surfaces.^19^ Expanding on this knowledge, the current study specifically focuses on exploring the relationship between microstructure density and binding capacity. Notably, a significant increase in molecular binding capacity is observed by reducing the center-to-center distance between pillars from 197 µm to 109 µm.

Moreover, this research sheds light on the potential of micropillar arrays as reliable platforms for biochemical experiments by integrating CRISPR-Cas assays. Specifically, by employing the designed FRET-CRISPR assay, channels with higher micropillar density exhibit enhanced detection sensitivity compared to those with lower micropillar density. This finding highlights the capacity of microstructures to augment the sensitivity of CRISPR-based assays. Future improvements in detection sensitivity can be pursued by exploring micropillars with higher aspect ratios.^29, 30^ Recent studies have revealed that the edges of micropillars can contribute to increased surface molecular capacity^31^, presenting another avenue to enhance the detection sensitivity of CRISPR-Cas assays. In addition, the rapid development of microscale additive manufacturing techniques could potentially further increase the detection sensitivity by incorporating multi-scale hierarchical structures.^32, 33^

This study makes a significant contribution by evaluating two chemical modification approaches for PDMS surfaces. The approaches involve treating the surfaces with APTMS alone or APTMS and glutaraldehyde. Our results show that both treatments have comparable molecular binding capacities. However, hen coupled with FRET assays, which involve quenching the fluorescence intensities of immobilized QDs using attached quencher probes, the surface treated with APTMS alone exhibits a higher fluorescence difference before and after quenching and the treatment protocol is simpler without using glutaraldehyde. This finding not only enhances the current study but also provides valuable insights that can guide and inform future studies and applications involving FRET-based assays.^35, 3034^

Another significant contribution of this study is the development of a valuable method to prevent bubble formation in PDMS channels. PDMS is known for its highly hydrophobic nature, which often leads to the formation of bubbles when fluids are introduced into the channels.^36^ These bubbles can get trapped inside a microfluidic device, potentially causing dis-uniform in downstream biochemical reactions. To address this issue, a simple and effective approach was implemented in this research. The PDMS channels were subjected to vacuuming and then immersed in water, allowing the trapped air inside the channels to be withdrawn by filling the channels with water. Unlike some published methods that require specific chemical coatings to prevent bubble formation^37, 38^, which may interfere with downstream biochemical reactions, our approach is versatile, straightforward, and compatible with a wide range of biochemical reactions conducted within the channels.^39, 40^

Human papillomavirus is responsible for over 90% of cervical cancers, making it the fourth deadliest cancer in women and the most prevalent pathogen associated with female cancers.^41^ In our study, we specifically targeted the HPV 16 fragment for detection due to its significant carcinogenic potential, being present in approximately 50% of cervical cancer cases.^42^ To ensure the specificity of the FRET-CRISPR detection method developed in this work, we utilized the HPV 18 fragment as the negative control. Although both HPV 16 and HPV 18 are high-risk HPV types, they differ in terms of cancer histological types and clinical management.^43, 44^ Thus, employing HPV 18 as the negative control enabled us to simulate clinical scenario to discriminate between these two high-risk HPV types. Since the guide RNA can be precisely programmed to identify specific target sequences, our approach can be expanded to encompass the detection of diverse pathogens exhibiting comparable strains.

In this study, ten different micropillar channels were actuated using a multi-syringe pump, allowing for the simultaneous execution of multiplex reactions. To address the growing demand for multiplex detection in clinical settings, the current single-layer PDMS channel design can be upgraded to a double-layer channel configuration, incorporating an additional pneumatic control layer.^45, 46^ This modification facilitates precise control of multiple fluidic flows simultaneously, thereby enhancing automation and scalability. Previous research has already demonstrated the effectiveness of a similar design in handling up to 80 samples in parallel.^47, 48^ Furthermore, this study demonstrated the application of the micropillar channel for the detection of dsDNA fragments of HPV 16 virus using the CRISPR-Cas12a assay. In fact, the versatility of the micropillar channel extends beyond this specific assay. By employing alternative assays such as CRISPR-Cas13, the channel can also be utilized for the detection of viral RNA sequences.^49, 50^

## Materials and Methods

### a. Preparation of the HPV fragment

The L1 gene fragment sequences of human papillomavirus (HPV) types 16 and 18 were incorporated into the pCDNA3.1(+) plasmid by Genscript. To amplify the specific regions of interest, a TwistAmp® Basic kit from TwistDx was employed following the recommended procedures. After an incubation at 37°C for 15 min, the amplicons underwent subsequent analysis and purification steps. Verification of the amplified fragments was conducted through DNA sequencing (Poochon Scientific), along with the use of electrophoresis. For precise quantification of the electrophoresis bands, the Gene Clean II method by MPbio was employed for efficient purification. The purified fragments were then subjected to re-quantitation using spectroscopy.

### b. Device fabrication

To fabricate high aspect-ratio microstructures, laser micromachining (Oxford Lasers) was employed to create microholes with a diameter of 25 µm at the center of a 5 mm x 10 mm tungsten foil. Following the drilling process, the foil was cleaned with ethanol, dried with nitrogen, and silanized overnight in a vacuum desiccator using a silanization solution (Sigma-Aldrich, #85126). The polydimethylsiloxane (PDMS) pre-polymer (SYLGARD 184 Silicone Elastomer Base) was thoroughly mixed with the SYLGARD 184 Silicone Curing Agent with a ratio of 10:1 and subsequently poured onto the silanized foil placed in a petri dish. To eliminate any trapped air bubbles, the mixture was degassed using a desiccator and cured by incubating in an oven at 65 °C for 2 h. Once fully cured, the PDMS slab was peeled off from the foil and 1 mm diameter holes were punched at both ends of the PDMS. Next, both the punched PDMS and a glass substrate were cleaned with ethanol and deionized water and exposed to oxygen plasma (Electro-Technic Products) for 45 s before being adhered together to form a channel. The channel was then immediately baked on a hotplate at 125 °C overnight.

### c. Surface modification

To ensure accurate surface modification within the channels, a thorough degassing process was performed to prevent the formation of bubbles during reagent injection. The channel was degassed using a desiccator for 15 min, followed by immersion in nuclease-free water. Subsequently, the trapped air was withdrawn, and the channel was filled with water, thereby creating an optimal environment for subsequent processes. The channel was injected with a freshly prepared solution of 10% (3-Aminopropyl)trimethoxysilane (APTMS, Sigma-Aldrich, #281778) in ethanol and incubated for 10 min. Subsequently, any excess untreated APTMS was eliminated by flushing the channel with 100 µL of ethanol at a flow rate of 25 µL/min, utilizing a syringe pump (WPI, #SP2201). The channel was then placed on a hot plate at 125 °C for 10 min.

Following the heating step, streptavidin-coated QDs (ThermoFisher Scientific, #Q10001MP) were injected into the channel and allowed to incubate for 5 min. To ensure the removal of any unbound quantum dots, a thorough rinsing of the channel was carried out using 200 µL of nuclease-free water, actuated by the syringe pump.

### d. CRISPR-Cas12a-based HPV detection

Firstly, a preassembly step was conducted by incubating AsCas12a (IDT, Inc, #10001272) at a concentration of 50 nM with 62.5 nM crRNA at room temperature for 10 min. Next, the AsCas12a-crRNA complexes were combined with 1 µL of binding buffer (New England BioLabs, #B6002S), 4.5 µL of biotinylated quencher probes, and 3.625 µL of nuclease-free water. Following this, 2 µL of target DNA was added, initiating the CRISPR reaction, and allowed to incubate at 37 °C for a duration of 2 h.^6, 51^

### e. Isolation of un-cleaved quencher probes though the IMPACT

Following the target identification and trans-cleavage by CRISPR, the resulting product with a volume of 15 µL was introduced into the surface modified channel and allowed to incubate at room temperature for 20 min. Subsequently, to remove any unbound probes and CRISPR-Cas complexes, the channel was rinsed with 200 µL of nuclease-free water at a flow rate of 25 µL/min.

### f. Fluorescence quantification

The channels were imaged using an inverted microscope (Zeiss 880), and the fluorescence intensity was measured using ImageJ software. Each channel was photographed five times, with randomly selected fields of view for each capture. To evaluate the detection performance, the fluorescence intensity of each channel was measured both before and after the CRISPR cleavage and the intensity difference was utilized to assess the sensitivity and specificity.

## Acknowledgements

This study was supported by The National Institutes of Health R35GM 142763.

